# Indigenous knowledge of plant uses by the community of Batiaghata, Khulna, Bangladesh

**DOI:** 10.1101/2020.07.22.216689

**Authors:** Tama Ray, Md. Sharif Hasan Limon, Md. Sajjad Hossain Tuhin, Arifa Sharmin

## Abstract

Southwestern region of Bangladesh is very rich in floral diversity, and their diversified uses. An extensive survey was conducted to investigate ethnobotanical applications of botanical species by the community of Khulna, Bangladesh. We focused on plants and community relationships, identify the most important species used, determine the relative importance of the species surveyed and calculated the Fidelity level (FI) and Cultural Significance Index (CSI) concerning individual species. In total, we have listed 136 species of 114 genera under 52 families, of which 32% (45 species) were used for folk medicine. Inheritance of traditional knowledge of medicinal plants was the primary source of knowledge acquisition through oral transmission over the generations. However, only 34% of the informants were traditional herbal practitioners. Most of the medicinal uses are primly associated with anti-inflammatory, anti-microbial, antiseptic, expectorant, antidote, fever reduction, and pain relief.

## 1. Introduction

Geographically, most of the territories of Bangladesh are formed by a delta plain with a tropical monsoon climate. It is vibrant with a vast biological diversity (Chowdhury and Koike, 2010) and lies under the Indo-Burma biological hotspot area (Mukul *et al.*, 2008). In total, 7,000 floral species were listed from this area with several endemic plants, and 50% of them are herb, 35% shrub, and woody climber and 15 % tree (Rahaman, 2004). Meanwhile, angiosperms are dominated in checklists with 5,700 species followed by 1,700 pteridophytes, 500 medicinal plant species, 130 fiber yielding plants species, 68 woody legume species, 29 orchid species and three species of gymnosperm (Mukul *et al.*, 2008). However, Ahamed *et al.*, (2007-2009), has enlisted 3611 angiosperm, 195 pteridophyte species, and seven gymnosperm species, excluding Bacteria and fungi. In addition to that, there are 750–800 more tree species found in these areas, including indigenous, exotic, and naturalized ones (Irfanullah, 2011). Among all different regions, the hilly region is the richest one in consideration of floristic diversity and richness with 2,260 plant species (Mukul *et al.*, 2008).

In Bangladesh, humans and plants share the natural habitats with traditional bonding and influence each other (Partha, 2014). Still, at most of the aspects of biological and economic needs, people depend on plants for food, shelter, construction materials, clothing, medicines, rituals, fuelwood, household implements, musical instruments, pesticides and so on (Gemedo-Dalle *et al.*, 2005). This dependency became the grass-root basis of species conservation for humans (Singh *et al.*, 2002). But at present, people are using trees in an exploiting manner for the economy (Idu, 2009; Kargioglu *et al.*, 2008). As a result, number of plants species is decreasing at faster rate, along with the knowledge of traditional use of those species (Balick and Cox 1996; Avocèvou-Ayisso *et al.*, 2012). Several studies have identified scarcity of ethnobotanical information and the lack of transmission of ethnobotanical knowledge from generation to generation as the crucial factors in disappearance of species from a locality (Khan *et al.*, 2018). Currently, ethnobotanical knowledge about medicinal plants found exposure to the scientific community and several studies have been conducted regarding this issue (Yusuf *et al.*, 2006; Partha and Hossain 2007; Roy *et al.*, 2008; Rahmatullah *et al.*, 2012; Uddin *et al.*, 2012). But most of the research missing the other usages bear comparatively the same importance in conservation and site-specific parameters. So, we considered this in this study to identify species composition and diversity, analyze the uses of species and their mode of use, and evaluate the value or importance of species within the culture.

## 2. Methodology

### 2.1 Study Area

The study was conducted in Batiaghata Upazila under Khulna district. It is located between 22°46’07” N to 22°37’50” N and 89°24’14” E to 89°31’47” E (Fig. 1). Batiaghata experiences a subtropical climatic condition with a mild winter from October to March, hot and humid summer in March to June and moist, warm rainy monsoon in June to October. In December-January, the temperature fell to the lowest at 12-15°C, and it reached highest in April-June at 41 -45 °C. Most of the rainfall during June to October. July is the month of maximum precipitations with 20-25 days of rain. Average wind speed is over 8 Km/h during April-August, which is the highest value for this area (BBS, 2014).

**Figure 1:**
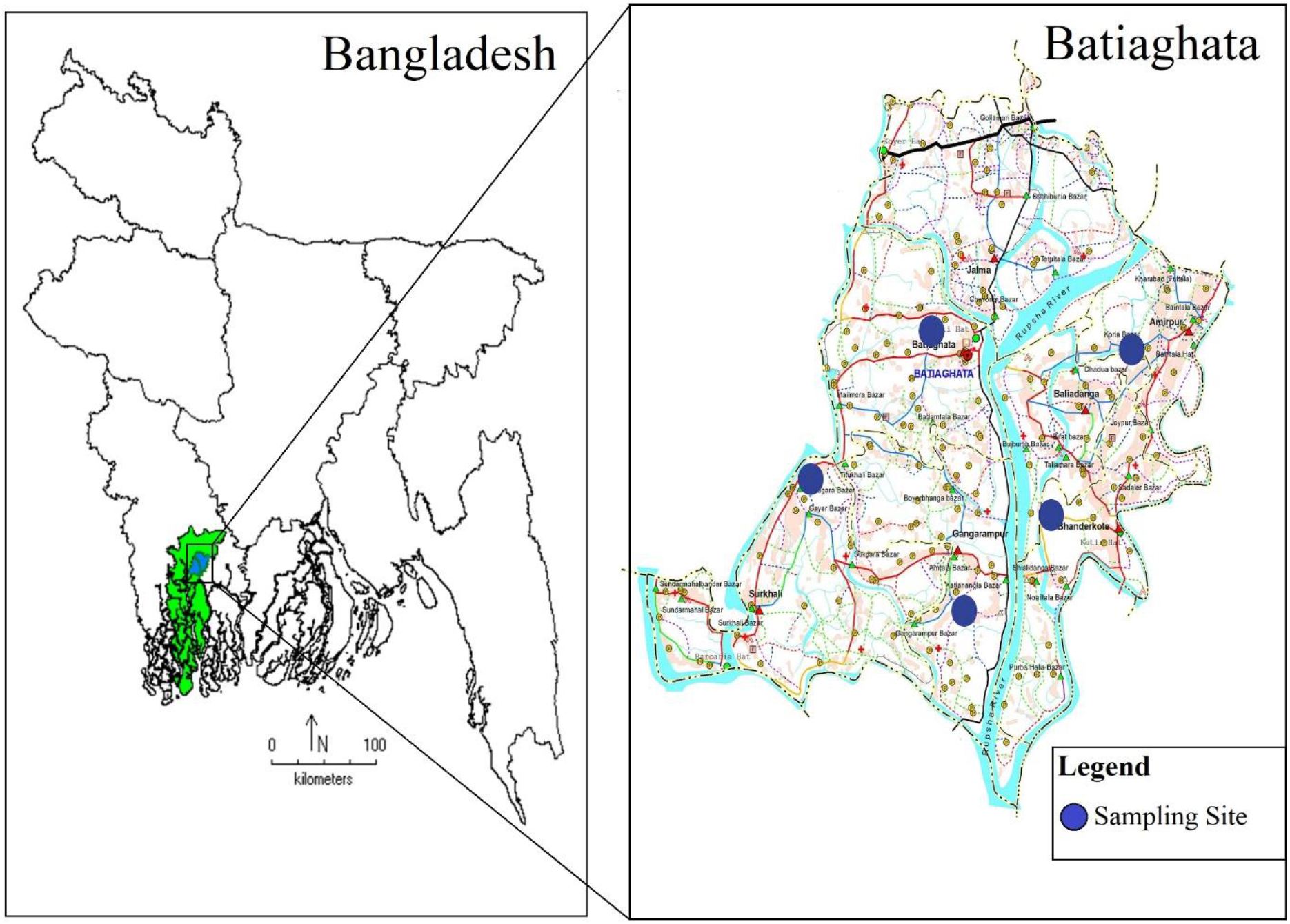
Map showing the study area.

### 2.2 Demography

Like other Upazilas of Bangladesh, Batiaghata is densely populated with a total population of 140,574 in which 72,717 are males and 67,857 females with 40,779 units of households. Among the total population, Muslims dominate with 79,301 along with 60,894 Hindu, 285 Buddhist, 6 Christian, and 85 others. Administratively, Batiaghata Upazila has seven unions named Amirpur Union, Baliadanga Union, Batiaghata Union, Bhandarkote Union, Gangarampur Union, Jalma Union, and Surkhali Union. Almost 91% of people are engaged in agriculture, followed by service (7%) and industry (2%) (BBS 2011, 2013).

### 2.3 Sampling Design

A reconnaissance survey was conducted in the study area before questionnaire preparation to obtain general information about the villages and the villagers. Depending on this survey, a semi-structured questionnaire was prepared for ethnobotanical information collection. Five unions (Batiaghata Union, Baliadanga Union, Gangarampur Union, Amirpur Union, and Surkhali Union) have been selected and one village from each union was chosen to survey by random selection. In total, 150 households were studied in this study, where 30 houses were chosen randomly from every village for data collection. Interviewees were divided into five age groups (20-34, 35-49, 50-64, 65-79, 80 and above) as age plays a distinctive role in ethnobotanical knowledge (Nawash *et al.*, 2014). Cited plant species were checked physically, photographed, and voucher specimens were collected for further identification and conservation. Collected specimens were analyzed and identified based on the key provided by Hooker (1872-1890), Prain (1903-04), Kanjilal *et al.*, (1934-1940), Deb (1983), Matthew (1999) and Ahmed et al. (2007-09).

### 2.4 Calculations

After collecting data, they were categorized according to their specific use such as food, medicine, construction, fuel, ornamental, and others and analyzed according to the following indices.

a. Fidelity level, FL = Ip − Iu * 100% (Friedman 1986; Hoffman and Gallaher 2007) Here, Ip = Number of informants who cited the species for the particular use. Iu = Total number of informants that mentioned the plant for any use.
b. Cultural significance index, CSI = ∑ (i*e*c) *CF (Turner 1988; Stoffl *et al*, 1990; Hoffman and Gallaher 2007) Here, I = species management where, 1 indicates non-managed and 2 indicates managed. E = Use preference where 1 indicates non- preferred and 2 indicates preferred. C = Use frequency where 1 indicates rarely used, and 2 indicates frequently used. CF (Correction factor) =Number of citations for a given species divided by the number of citations for the most mentioned species.

## 3. Result

In total, 136 species of 114 genera under 52 families have been identified throughout this study. Among the counted species, 41% species were utilized as food, followed by 30% medicine, 14% constructional timber, 11% ornamental, and 4% other uses (Fig. 2a). Forty-four species have been cited by the informants to be used in the treatment of various human diseases, including respiratory, digestive, liver, skin, rheumatism, diabetes, cancer, and other disorders. Among the 150 informants interviewed, 72.6% were males, and 27.4% were females, with 74.3% above 50 years of age. Inheritance of traditional knowledge of medicinal plants was the primary source of knowledge acquisition through oral transmission over the generations. However, only 34% of the informants were traditional herbal practitioners, with the remaining majority (66%) of informants having no professional practice of herbal medicine.

**Fig. 2:**
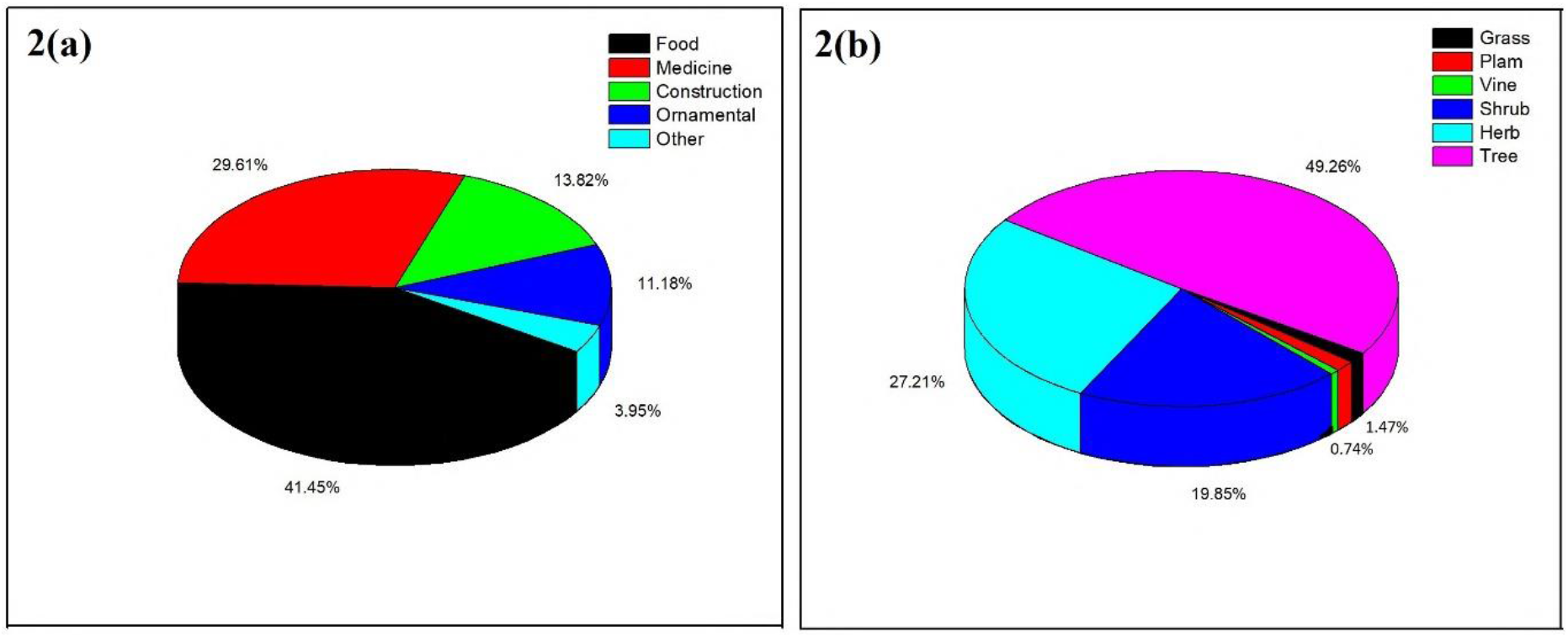
Composition of identified species, (a) different uses, (b) life forms.

One hundred thirty-six floral species have been cited in the homestead of the study area belonging to 52 families and 114 genera. Fabaceae (17) found the most dominant family followed by Anacardiaceae (6), Myrtaceae (6), Apocynaceae (5), Arecaceae (5), Malvaceae (5), Moraceae (5), Solanaceae (5) (Fig. 3). Horticultural species like Syzygium (5), Terminalia (4) Artocarpus (2) found most dominant over other species. The life forms and growth habits of plants were distributed into 49.3% trees, 19.9% herb, 1.5% grass, 1.5% palm, and 0.7% vine (Fig. 2b). Most of the medicinal uses are undoubtedly associated with anti-inflammatory, anti-microbial and antiseptic, antibacterial, expectorant, antidote, fever reduction, and pain relief.

**Fig. 3:**
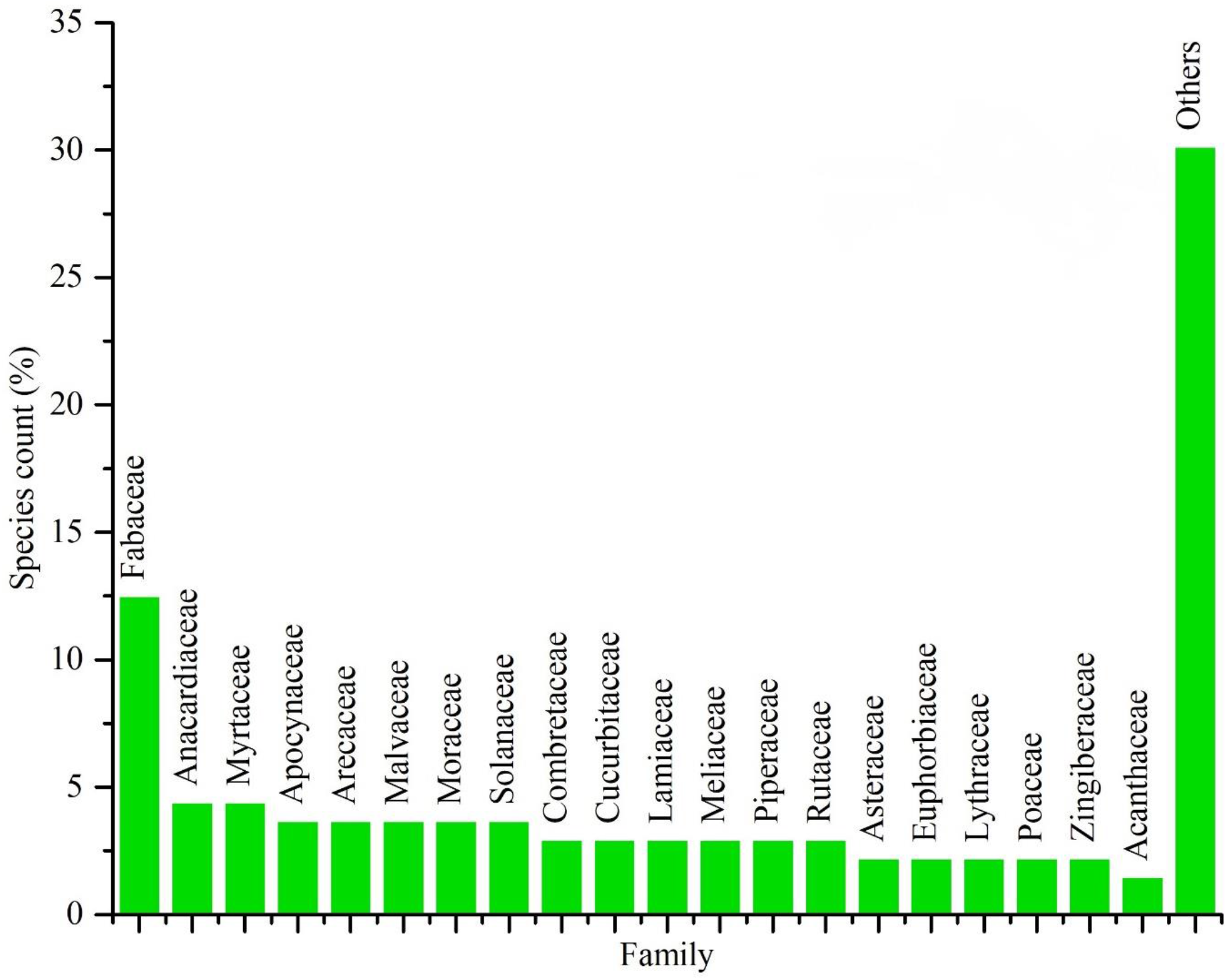
Dominant Families of the study area.

*Mangifera indica* found most cited species over the study area with a high informant consensus (IC) of 86, followed by *Areca catechu* (79), *Cocos nucifera* (76), *Ocimum tenuiflorum* (73); *Swietenia macrophylla* (73), *Albizia lebbeck* (69) and so on. In terms of Fidelity Index (FI %), 114 species scored 100%, which indicates the dedicated use of those species without any alternatives. Cultural Significance Index ranges from 13.58 to 0.08. Six species lie above ten, namely, *Ocimum tenuiflorum* (13.58), *Mangifera indica* (13), Areca catechu (11.94), *Cocos nucifera* (11.49), *Swietenia macrophylla* (11.03), *Albizia lebbeck* (10.43) (Table 1).

**Table 1.**
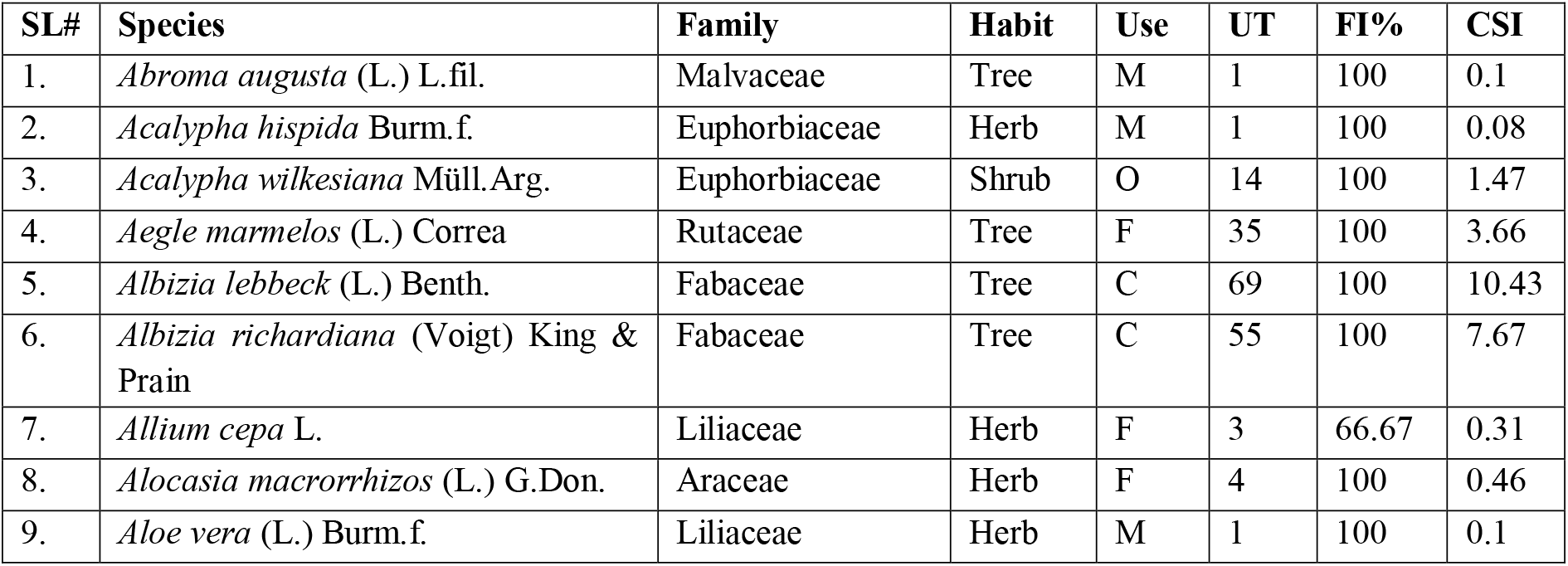
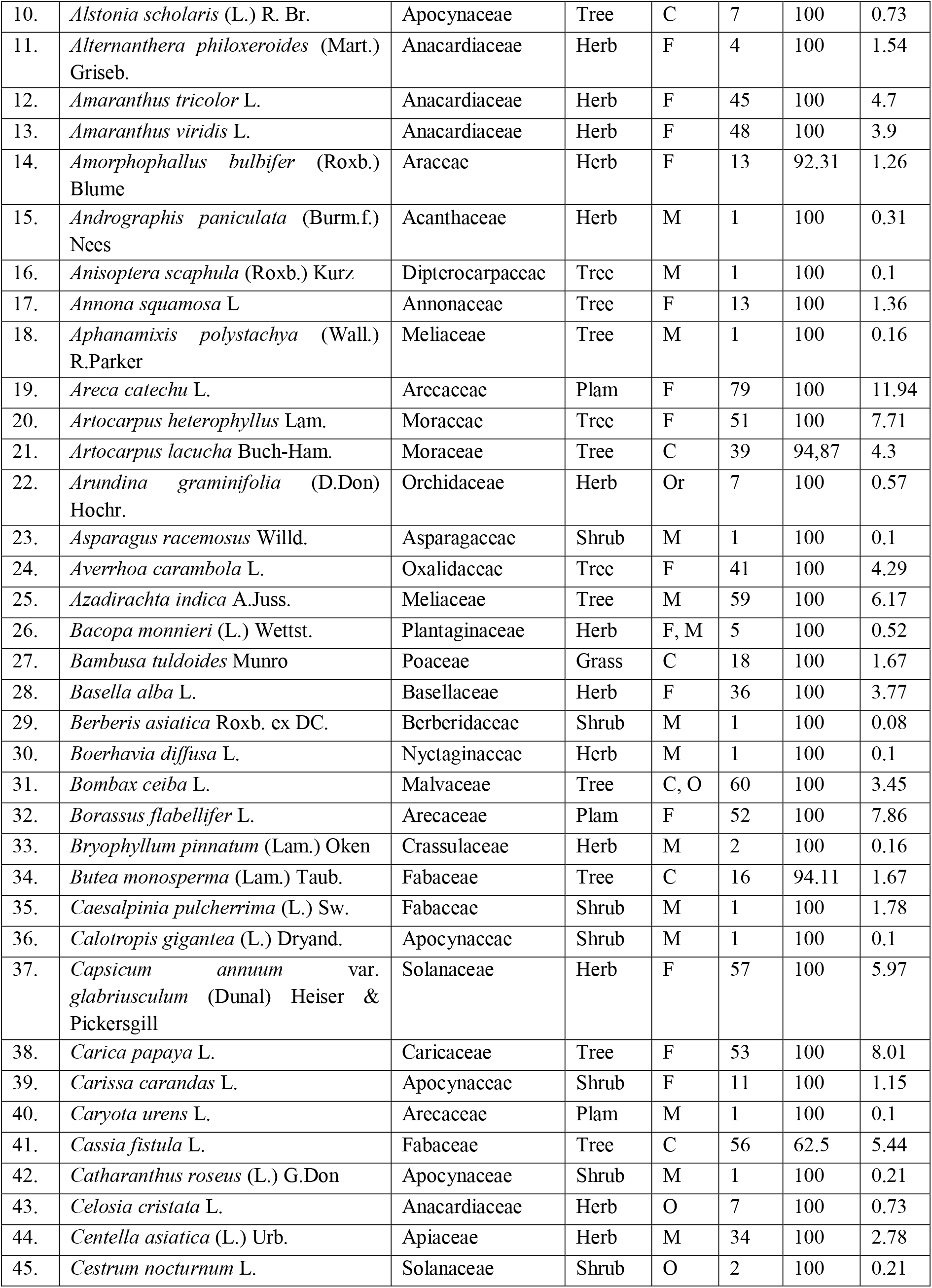
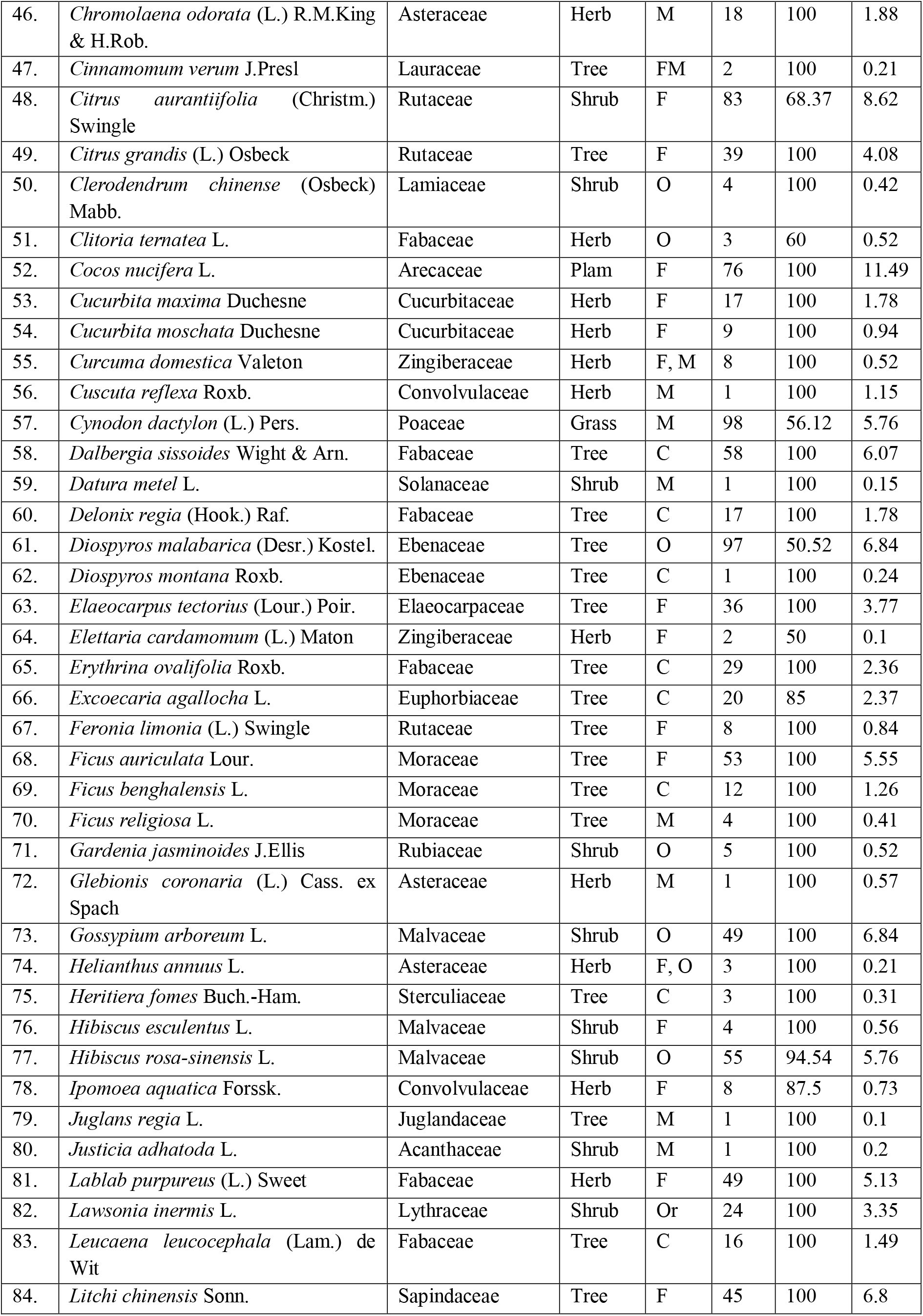
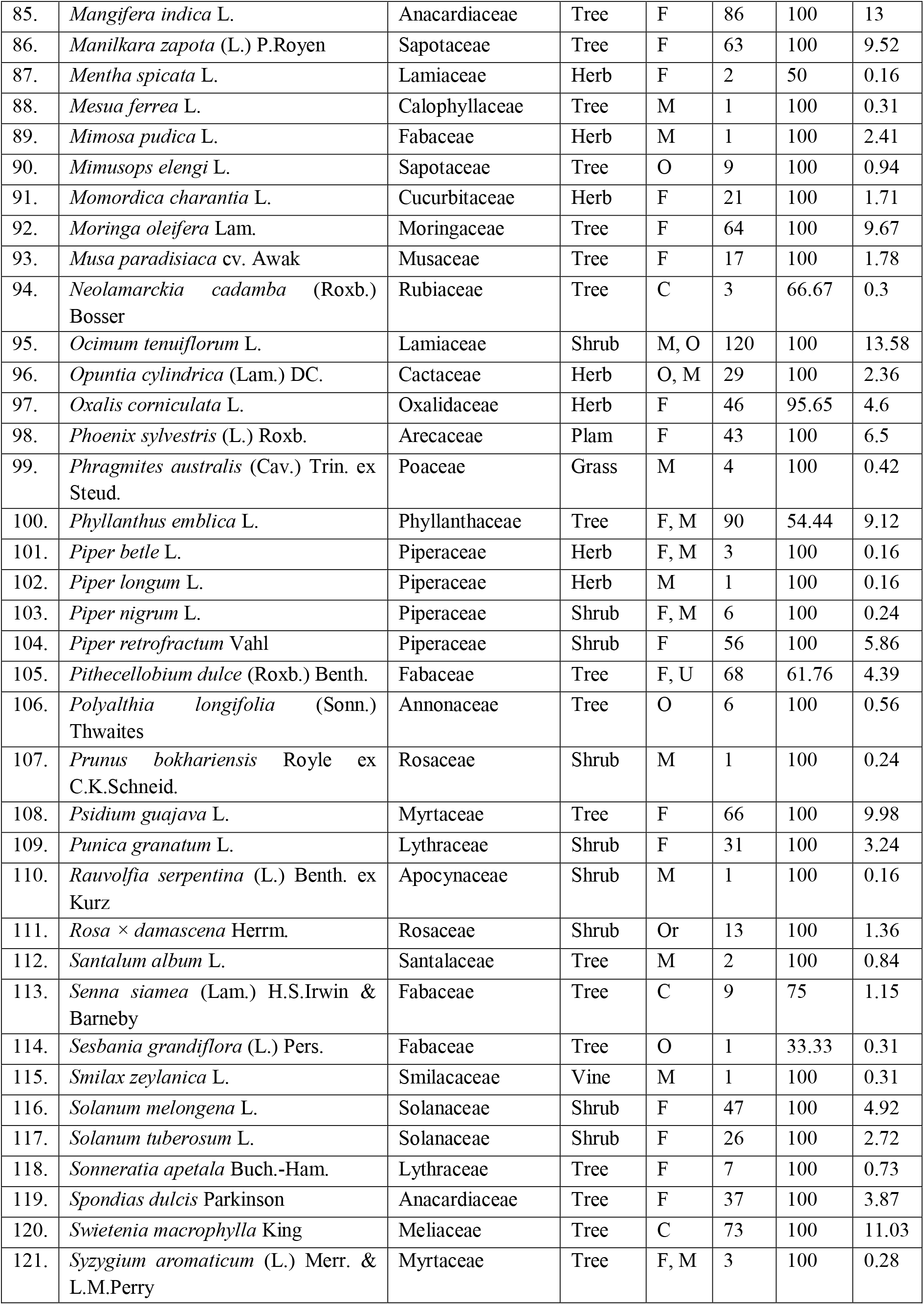
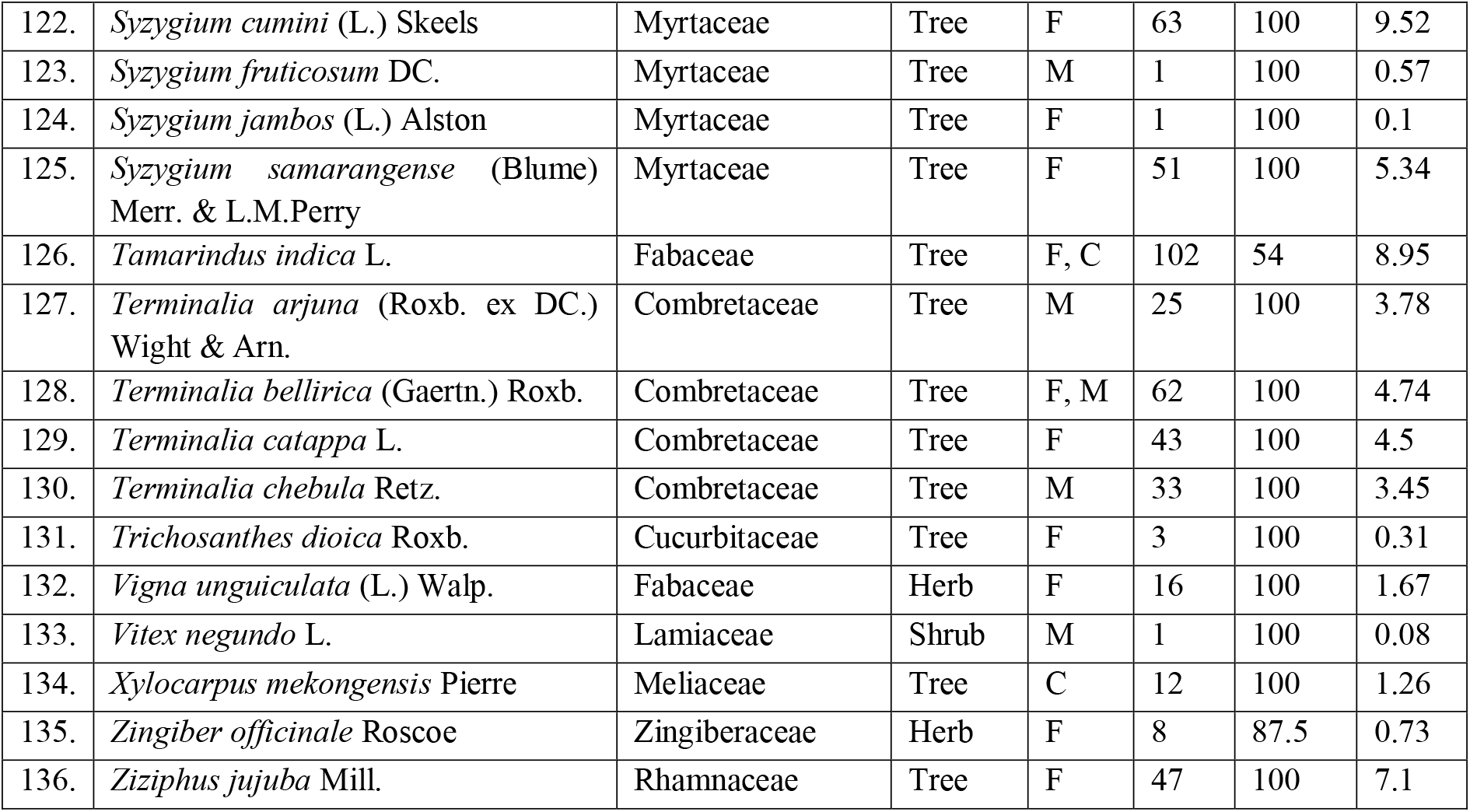
Summary of the study. Ethnobotanical indices of Batiaghata, Khulna (IC=Informant Citation; UT=Used Total; FI=Fidelity Level; CSI=Cultural Significance Index; F=Food; M=Medicine; F=Fuel; C=Construction; Or=Ornamental and O=Other).

## 4. Discussion

A considerable amount of floral species has been used traditionally by the local community with diversified implications in the study area. Very few studies have been done in this region to preserve the ethnobotanical knowledge inherited by generations. In 2009, Nawaz et al. listed 26 plant species from 22 families used in ethno-medicine from Khulna and Jessore, Mollik *et al.*, (2009) identified 33 species in folk medicine from Khulna division and Ray and Mandol (2018) describe 25 species from Shyamnagar, Satkhira near Sundarbans. However, at present studies, 136 species of 114 genera under 52 families were identified, of which 45 species have been reported to be used in folk medicine.

The use-value indicates the total number of uses of a specific species and two types of tally have been used for calculations, Uses Total (Researcher Tally) indicated specific applications and Use- Value indicated individual allocation. However, in researcher tally, the uses were recorded, ranked, and summed, which showed a similar contingency to previous studies (Rahman, 2013; Ray and Mandol, 2018; Faruque *et al.*, 2018). In this study, food, fuel, medicine, construction, ornamental, and other categories were used to investigate multiple uses of a single species and numerous species for individual use.

We have found the highest species use-value for *Mangifera indica* (0.95), and total use is also very high for it (86), which means the highest number of people engage with this species by its meaningful use at their daily life. Meanwhile, *Cuscuta reflexa* remains at the lowest level (0.01) for its minimal use by the lowest number of people from the participants. In terms of fidelity level, it describes the importance of the species for any specific purpose. It is used to identify the preferable species used by key informants for one particular treatment. Species having high fidelity levels is generally widely used for dedicated purposes. It also illustrates the number of informants in the percentage who state the use of certain species for the same purpose (Khan *et al.,* 2014). Following that, fidelity levels were calculated highest (100%) for most of the species (114), which represents the single specific use of those species. But Lower fidelity level indicates multiple uses of a species, and we found 24 species have various applications with lower fidelity levels. *Alocasia indica* has the fidelity level 100% having one primary purpose of use as food. 114 (83.33%) species have the highest (100%) fidelity level, which indicates single use of those species. *Sesbania grandiflora* found lowest FI% (33.33), indicating multiple uses (Table 01). Cultural Significant index (CSI) indicates the versatility of the application of a species along with the number of informants it uses, which means the spread of the use of the species (Prthiban *et al.,* 2016). We have calculated the highest CSI for *Ocimum tenuiflorum* (13.58), which indicates different uses like medicinal and worship purposes.

The identified species composition showed higher diversity in use and practice. Religion, social strata, and economic conditions plays pivotal role to regulate the level of applications and inheritance of ethnobotanical knowledge. The current study found that most of the information (66%) about ethnobotany was inherited through oral communications. The aged personals of the community have a significant role in transmitting this knowledge among the population. In addition to that, several professionals, locally named as Kabiraj or Gunim (traditional healer), also have a strong influence over folk medicine use at the community level. Local people also cited that the use and practice of folk medicine is diminishing. Limited use of this folk medicine also threatened the conservation of the species in the region. Currently, most of the medicinal purposes are limited to treat anti-inflammatory, anti-microbial and antiseptic, antibacterial, expectorant, antidote, fever reduction, and pain relief.

## 5. Acknowledgment

Authors are grateful to Late Md. Shirazul Islam Sheikh, Katianangla, Batiaghata, Khulna, for his active participation and supporting field data collection process.

